# A rapid and accurate approach for Prediction of interactomes from co-elution data (PrInCE)

**DOI:** 10.1101/152355

**Authors:** R. Greg Stacey, Michael A. Skinnider, Nichollas E. Scott, Leonard J. Foster

## Abstract

**Background:** An organism’s protein interactome, or complete network of protein-protein interactions, defines the protein complexes that drive cellular processes. Techniques for studying protein complexes have traditionally applied targeted strategies such as yeast two-hybrid or affinity purification-mass spectrometry to assess protein interactions. However, given the vast number of protein complexes, more scalable methods are necessary to accelerate interaction discovery and to construct whole interactomes. We recently developed a complementary technique based on the use of protein correlation profiling (PCP) and stable isotope labeling in amino acids in cell culture (SILAC) to assess chromatographic co-elution as evidence of interacting proteins. Importantly, PCP-SILAC is also capable of measuring protein interactions simultaneously under multiple biological conditions, allowing the detection of treatment-specific changes to an interactome. Given the uniqueness and high dimensionality of co-elution data, new tools are needed to compare protein elution profiles, control false discovery rates, and construct an accurate interactome.

**Results:** Here we describe a freely available bioinformatics pipeline, PrInCE, for the analysis of co-elution data. PrInCE is a modular, open-source library that is computationally inexpensive, able to use label and label-free data, and capable of detecting tens of thousands of protein-protein interactions. Using a machine learning approach, PrInCE offers greatly reduced run time, better performance, prediction of protein complexes, and greater ease of use over previous bioinformatics tools for co-elution data. PrInCE is implemented in Matlab (version R2015b). Source code and standalone executable programs for Windows and Mac OSX are available at https://github.com/fosterlab/PrInCE, where usage instructions can be found. An example dataset and output are also provided for testing purposes.

**Conclusions:** PrInCE is the first fast and easy-to-use data analysis pipeline that predicts interactomes and protein complexes from co-elution data. PrInCE allows researchers without bioinformatics proficiency to analyze high-throughput co-elution datasets.

## 1 Background

The association of proteins into complexes is common across all domains of life [1, 2]. Indeed, most proteins in well-studied proteomes are involved in at least one protein complex [3, 4]. Therefore, understanding the roles, mechanisms, and interplay of protein complexes is central to understanding life.

A proteome of 1,500 proteins has over one million possible binary protein-protein interactions (PPIs) and many more potential higher-order complexes. Because of this combinatorial explosion, even relatively simple proteomes can yield rich, complex interactomes. High-throughput or high-content methods that identify many PPIs simultaneously are therefore valuable to efficiently map these networks. There are currently three general methods for doing this: The first, yeast-2 hybrid (Y2H), operates by incorporating modified bait and prey proteins in a genetically modified yeast cell, such that a PPI between bait and prey drives transcription of a reporter gene. Affinity purification mass spectrometry (AP-MS), a second technique, involves immunoprecipitation of proteins of interest (baits) [5]. While powerful, both techniques face limitations. For one, tagging proteins, typically with Gal4 in the case of Y2H or an epitope-antibody combination for AP-MS, creates non-endogenous conditions that can disrupt protein binding sites and increase the number of false negatives.

The third general approach, collectively termed co-fractionation approaches, involves resolving complexes by either chromatography or electrophoresis and assigning interacting partners based on the similarity of fractionation profiles [6–8]. While there are similarities in how the data from these methods are treated, there are also unique considerations for each one. Being more established methods, Y2H and AP-MS have several excellent approaches for data analysis [5, 9, 10]. However, there does not yet exist a gold standard tool for analyzing co-fractionation data. We [11] and others have previously reported pipelines for analyzing co-fractionation data, although existing approaches use other external sources of data, e.g. co-evolution, in addition to co-fractionation data [6, 12]. Optimally though, an interactome should be derived from co-fractionation data alone, using other data only for benchmarking. To this end, here we describe an open-source pipeline for analyzing co-fractionation data: PrInCE (Prediction of Interactomes from Co-Elution). PrInCE represents a major conceptual advance over preliminary bioinformatics treatments published by our lab, which provided basic data extraction and curve fitting tools for co-elution data [8, 11]. Improvements include ranked interactions, improved user interface, and extensive documentation. Importantly, PrInCE uses machine learning methods which greatly improve its performance. We benchmarked the performance of PrInCE versus a previous version [11] and demonstrate a 1.5-to-2-fold improvement in the number of predicted PPIs at a given false disovery rate with a 97% decrease in computational cost. This pipeline is freely available for download [13].

## 2 Methods

### Pipeline overview

The workflow of the pipeline is divided into five modules: 1) identification of Gaussian-like peaks in the co-fractionation profiles (*GaussBuild*.m); 2) correction for slight differences in the separation dimension between replicates (*Alignment.m*); 3) comparison of differences in protein amounts, i.e. fold changes, between conditions (*FoldChange.m*); 4) prediction of PPIs within each condition (*Interactions. m*); and 5) construction of protein complexes from the predicted PPIs (*Complexes.m*). The first two modules, i.e. *GaussBuild.m* and *Alignment.m*, are pre-processing steps, while the remaining three modules compute protein abundance changes and predict protein interactions and complexes (Figure 1).

**Figure 1:**
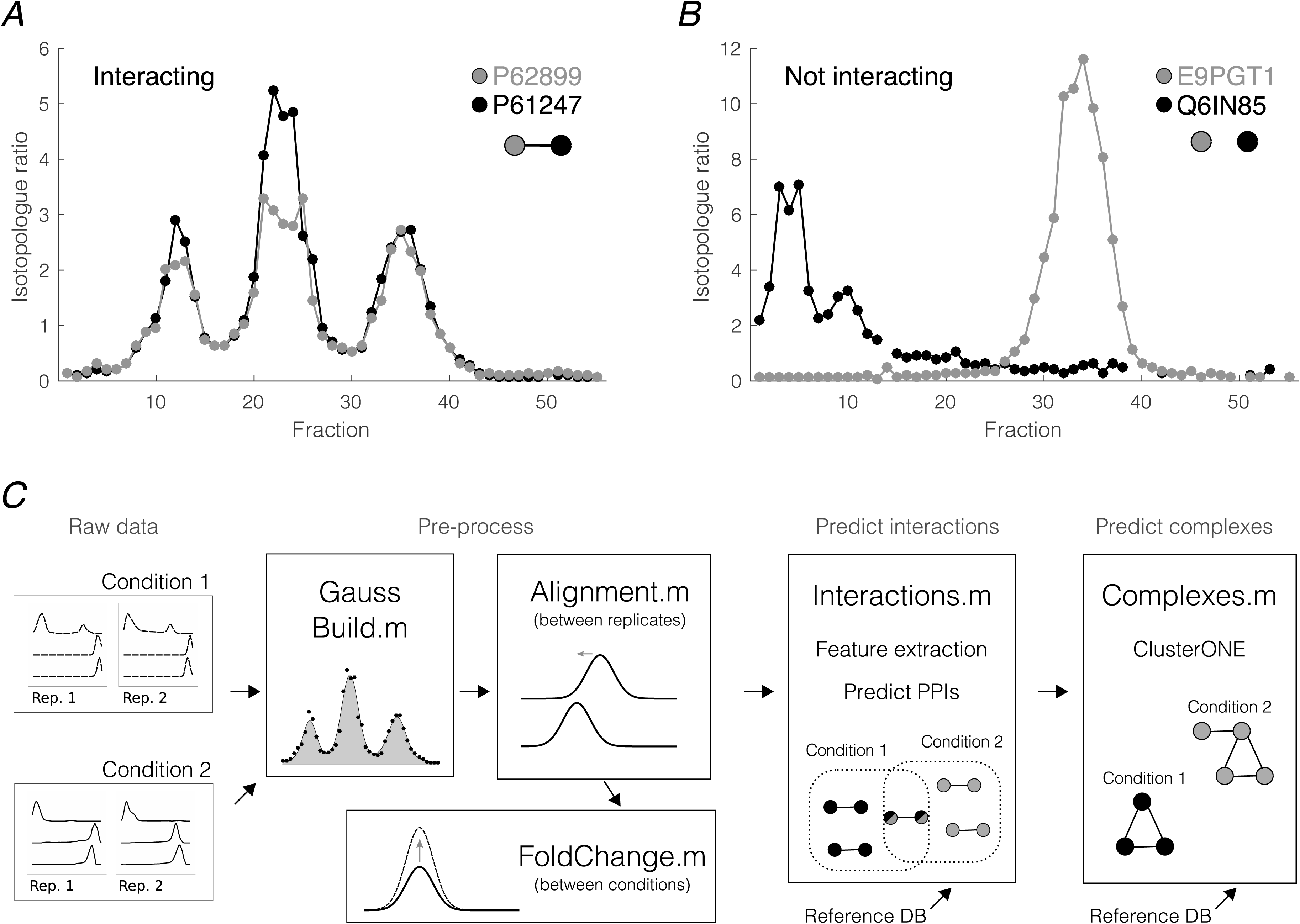
Pipeline overview. A. Co-fractionation profiles from known interactors, ribosomal proteins P61247 (black) and P62899 (grey). B. Co-fractionation profiles from non-interacting protein pair, Q6IN85 (black) and E9PGT1 (grey). C. Pipeline workflow. Raw data consists of co-fractionation profiles grouped by replicate and condition. In pre-processing, Gaussian mixture models are fit to each co-fractionation profile to obtain peak height, width, and center. If there are multiple replicates, the Alignment module adjusts profiles such that Gaussian peaks for the same protein occur in the same fraction across replicates. Changes in protein amounts between conditions, i.e. fold changes, are computed in the FoldChange module. Inter-actions between pairs of proteins are predicted by first calculating distance measures between each pair of proteins and feeding these into a Naive Bayes supervised learning classifier. Known (non-)interactions from a reference database, e.g. CORUM, are used for training. Finally, the list of predicted pairwise interactions is processed by an optimized ClusterONE algorithm (Nepusz *et al.*, 2012) to predict protein complexes.

### Requirements

#### Software and hardware

PrInCE is available as a standalone program for Windows or Mac OSX, as well as a Matlab package. Matlab is not required to run standalone versions of PrInCE but it was selected initially due to superior curve fitting tools compared to other environments. After downloading and saving to a dedicated folder containing co-elution data, standalone PrInCE is directly accessed through its own icon. PrInCE can be downloaded for free [13]. Detailed documentation of all the code as well as further instructions for running the software are provided.

#### Datasets

This pipeline requires co-fractionation profiles of single proteins, where co-elution is evidence of co-complex membership. Each co-fractionation profile, e.g. a chromatogram, is a row in a *. csv* file. Co-fractionation profiles are grouped by both experimental condition and replicate number. Separate *.csv* files are used for different experimental conditions, and the replicate number of each chromatogram is recorded by a column in each file. We provide a test dataset on Github as an example of correct formatting.

#### Reference database of known complexes

This pipeline requires a reference database of known protein complexes. A portion of the proteins in these reference complexes must also be quantified in the experimental data, as the reference complexes provide the template by which novel interactions are predicted. We found that manually curated databases that rely on experimental evidence, such as CORUM [14], lead to a high number of predicted interactions.

## Pipeline workflow

### Data pre-processing (GaussBuild.m, Alignment.m)

Module *GaussBuild.m* uses Gaussian model fitting to identify the location, width, and height of peaks in the co-fractionation data. Any co-fractionation profile with data in at least five fractions is chosen for model fitting. First, single missing values in co-fractionation profiles are imputed as the mean of neighbouring data points. Remaining missing values are imputed as zeros, and co-fractionation profiles are smoothed by a sliding average with a width of 5 data points. Five Gaussian mixture models are fit to each profile. These models are mixtures of 1, 2, 3, 4 or 5 Guassians, respectively. Fitted parameters *A*, *μ*, and σ are the Gaussian height, center, and width, respectively. In order to reduce the sensitivity to outliers, robust fitting is performed using the *L*_1_ norm. For each profile, model selection is performed by selecting minimum AIC values.

Slight differences between the elution time of replicates are corrected by module *Alignment.m*, using the assumption that proteins with a single, well-defined chromatogram peak should elute in the same fraction in every replicate [11].

### Fold changes between conditions (FoldChanges.m)

Within a single replicate, the protein abundance ratio, i.e. fold change, is calculated between conditions for each protein (*FoldChanges.m*). If there are multiple replicates, this module also calculates significance using a paired t-test. Fold changes are calculated using data centered on the Gaussian peaks identified by *GaussBuild.m* [11].

### Predicting interactions (Interactions.m)

#### Quantifying co-fractionation with distance measures

PPI prediction begins by calculating the effective distance between the co-fractionation profiles of every pair of proteins. We use five distance measures to quantify different aspects of co-fractionation profile similarity. For all distance measures, a value close to zero signals high similarity between co-fractionation profiles. These five metrics are not exhaustive, but in practice we found there was little value in additional measures. For a pair of co-fractionation profiles *ci*, *cj*, these distance measures are

- One minus correlation coefficient, 1 − *R_corr_*: One minus the Pearson correlation coefficient between *ci* and *cj*.
- Correlation p-value, *pcorr*: Corresponding p-value to 1 − *R_corr_*.
- Euclidean distance between co-fractionation profiles *ci* and *cj*, *E*.
- Peak location, *P*: Calculated as the difference, in fractions, between the locations of the maximum values of *ci* and *cj*.
- Co-apex score, *CA*: Euclidean distance between the closest (*μ, σ*) pairs, where *μ* and *σ* are Gaussian parameters fitted to *ci and cj*. For example, if *ci* is fit by two Gaussians with (*μ, σ*) equal to (5, 1) and (45, 3), and *cj* is fit by one Gaussian with parameters (45, 2), 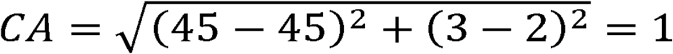. Thus chromatograms with at least one pair of similar Gaussian peaks will have a low (similar) Co-apex score.

#### Predicting interactions via similarity to reference

Combined with a reference database such as CORUM, these five distance measures can be used to predict novel PPIs. Our pipeline uses a machine learning classifier to do this [6, 15]. Specifically, we train a Naïve Bayes classifier, which evaluates how closely the distance measures for a candidate protein-protein pair resemble the distance measures observed for reference interactions. Distance measures are normalized such that their means are 0 and standard deviations 1. To reject uninformative distance measures, feature selection is performed prior to classification using a Fisher ratio > 2. The contribution of each feature to prediction performance depends on the dataset, although in general the most-informative (least-rejected) features are 1-*R_corr_*, *P*, and *CA*. Distance measures are combined across replicates (but not conditions) for each protein-protein pair. Class labels are assigned based on the reference database. Reference protein pairs that occur in the same complex are gold standard interactions (interacting or “intra-complex” label). Proteins that are found in the reference database individually but do not occur within the same complex are labeled non-interacting (“inter-complex”) and are false positive interactions [6]. Novel interactions are those where one or both members are not in the reference database.

The Naïve Bayes classifier returns the probability that putative protein pairs are interacting. Interaction probabilities are calculated separately for each experimental condition. We use a *k*-fold cross-validation scheme to avoid over-fitting. *k* = 15 is used as a tradeoff between computation time and classification accuracy. The classifier calculates an interaction probability for every protein pair. Self-interactions are not considered.

By applying a threshold to interaction probability returned by the classifier, protein pairs are separated into predicted interactions and predicted non-interactions. The probability threshold is chosen so that the resulting interaction list has a desired ratio of true positives (intra-complex) and false positives (inter-complex), quantified as precision *TP/*(*TP* + *FP*), where *TP* and *FP* are the number of true positives and false positives. The desired precision is chosen by the user.

Finally, we express the confidence of each predicted interaction by reformulating interaction probability as an *interaction score*. A predicted interaction’s score is equal to the precision of all predicted interactions with an interaction probability greater than or equal to it. Although interaction probability and score are largely equivalent, interaction score has two advantages. First, interaction score is more human readable, since the dynamic range of predicted interaction probabilities is often quite small. Second, the use of interaction score makes it trivial to generate interaction lists with a desired precision.

### Predicting complexes (Complexes.m)

Complexes are predicted from the list of pairwise interactions using the ClusterONE algorithm [16]. The primary benefit of ClusterONE over other algorithms is that ClusterONE can predict the same protein in multiple complexes. Two parameters, *p* and *dens* are optimized via grid search to produce the most reference-like complexes. *p* represents the number of unknown pairwise interactions, and *dens* is a threshold for the minimum density of a complex, where complex density is defined as the sum of weighted internal edges divided by *N*(*N*−1)/2. Parameters are optimized to maximize either the matching ratio [ 16] or geometric accuracy [17] between predicted and reference complexes. Since there are possibly multiple interaction lists – a list of all predicted interactions as well as lists specific to each experimental condition – complexes can be built for each experimental condition separately, as well as an overall complex set from the aggregate interactome.

### Test datasets

For this study, we tested PrInCE on four co-fractionation datasets, each composed of thousands of co-fractionation profiles (Table 1). D1, D2, and D4 were collected for recently published PCPSILAC experiments (D1 [18], D2 [11], D4 [8]). D3 is the raw intensity values of the medium channel of D1, which we included as a surrogate for non-SILAC data, and label-free data more generally.

**Table 1.**
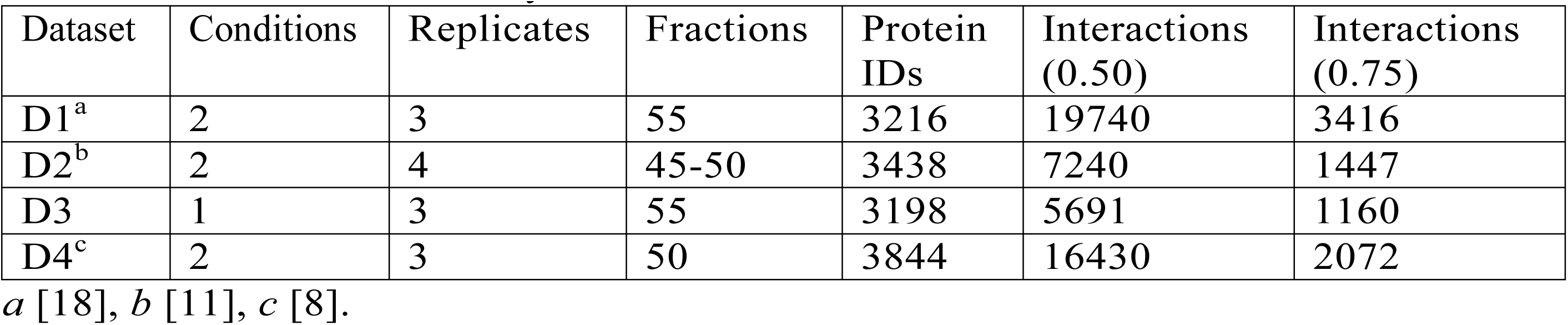
Test dataset summary

| Dataset | Conditions | Replicates | Fractions | Protein IDs | Interactions (0.50) | Interactions (0.75) |
| --- | --- | --- | --- | --- | --- | --- |
| D1<sup>a</sup> | 2 | 3 | 55 | 3216 | 19740 | 3416 |
| D2<sup>b</sup> | 2 | 4 | 45-50 | 3438 | 7240 | 1447 |
| D3 | 1 | 3 | 55 | 3198 | 5691 | 1160 |
| D4<sup>c</sup> | 2 | 3 | 50 | 3844 | 16430 | 2072 |

<sup>a</sup> [18], <sup>b</sup> [11], <sup>c</sup> [8].

### Validation of PrInCE output

Using these four datasets, we performed computational validations of PrInCE output. First, we tested whether our metric for ranking predicted interactions (interaction score) is consistent with other known evidence for protein interaction. To do so, we calculated the Spearman correlation coefficient between interaction score and these four other, independent measures of protein interaction: (i) whether protein pairs shared at least one Gene Ontology term within GO slim, a condensed version of the full GO ontology [19, 20]; (ii) the Pearson correlation coefficient of protein abundance across 30 human tissues, as taken from the Human Proteome Map (http://www.humanproteomemap.org/, [21]); (iii) whether protein pairs shared at least one subcellular localization annotation within the Human Protein Atlas Database [22]; and (iv) whether protein pairs shared a structurally resolved domain-domain interface, as identified by the database of three-dimensional interacting domains (3did) [23]. This validation was performed on predicted interaction lists with an interaction score of 0.50 or greater.

Second, we investigated whether predicted interactions were enriched over non-interactions for the same four measures (shared GO terms, tissue-dependent proteome abundance correlation, shared subcellular localization terms, and shared structurally resolved interfaces). For these interacting versus non-interacting enrichment analyses, we imposed a 10% breadth cutoff on all annotation terms, such that only annotation terms common to less than 10% of all proteins in the sample were used. As in [24], we also used the Jaccard index between protein pairs to quantify the extent of shared annotation terms across the entire Gene Ontology. This validation was performed on high-quality interaction lists (interaction score 0.75 or greater).

Third, we re-estimated the precision of our predicted interaction lists using an independent, previously described method [25]. Our definition of false positives as “inter-complex interactions” likely overestimates the number of false positives. To quantify the magnitude of this overestimation, we added random interactions between non-interacting proteins within the reference set to bring the average expression correlation coefficient of all interacting proteins within the reference dataset to the same level as in the predicted interactome under investigation. To avoid training and testing on the same reference interactions, we randomly withheld 1/3 of CORUM complexes as a validation set, and used the remaining 2/3 as a training set to train the Naive Bayes classifier and predict interactions. The average Pearson correlation coefficient in tissue proteome abundance was calculated for the resulting predicted interactions, and it was compared to interactions from the 1/3 of CORUM withheld for testing. We bootstrapped this procedure 100 times to re-estimate the precision of the protein interaction network.

Finally, following the network analysis of [24], we explored the topological properties of the predicted subgraphs by sequentially removing interactions under one of three schemes: (i) highest interaction score first, (ii) lowest interaction score first, or (iii) randomly. This analysis tests whether the interaction network consists of cores of tightly connected proteins linked by weaker or more spurious connections. If this is the case, removing weakest interactions first will fragment the network, increasing the number of unconnected subgraphs and lowering their average size, whereas removing the highest scoring interactions first will not fragment the network.

## 3 Results

PrInCE uses a machine learning approach to predict conditional interactomes from co-fractionation data. Four datasets were used to benchmark PrInCE versus a previous pipeline [11], which showed that PRInCE can discover twice the number of predicted PPIs (Fig. 2A) in less than one tenth the time (Fig. 2B). This improved runtime also includes the complex-building module, *Complexes.m*, that was not present in the previous version.

**Figure 2:**
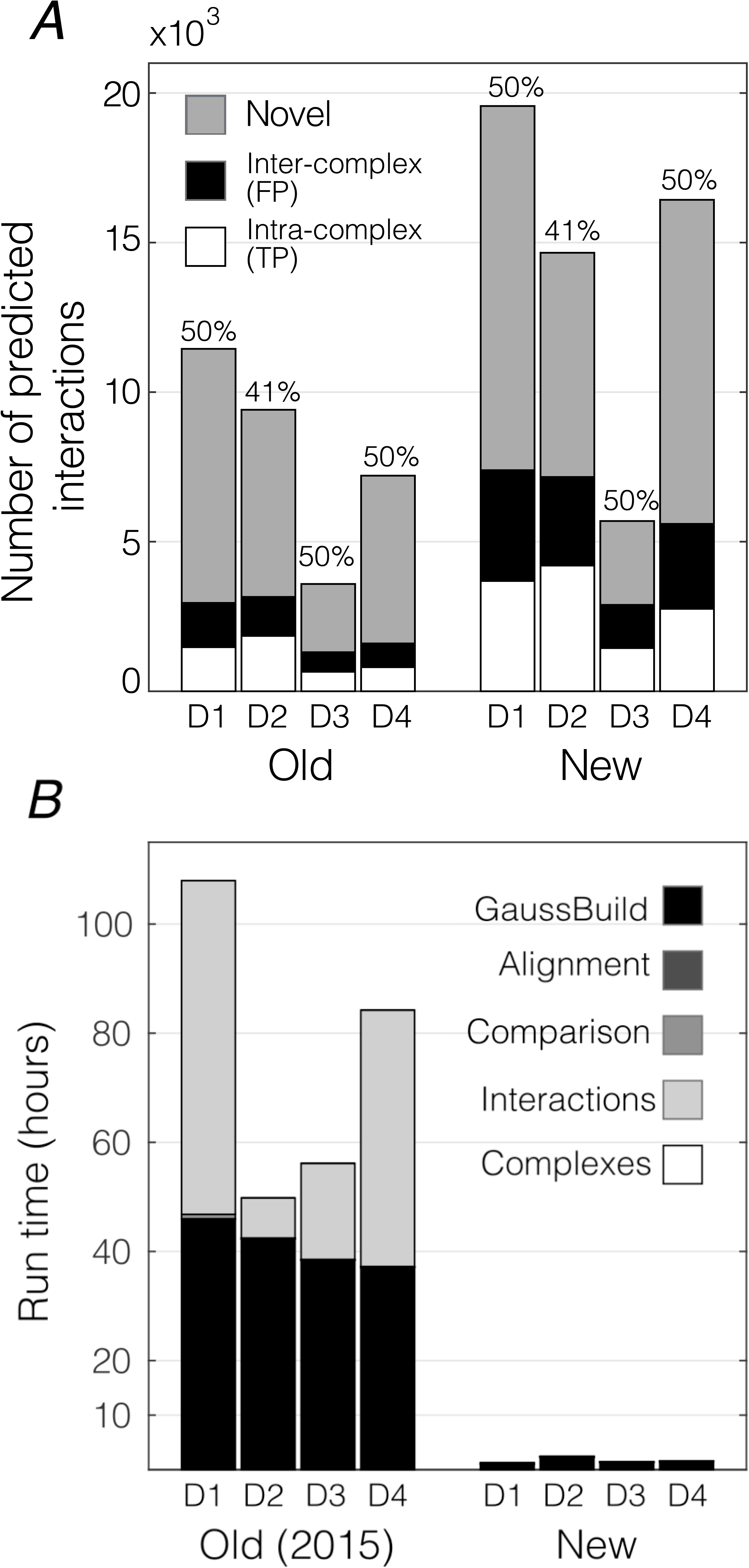
Improvements to predictive power and run time. A. Number of interactions predicted at 50% (D1, D3, D4) or 41% precision (D2). For previously published datasets (D1, D2, D4), precision values and interaction numbers reflect published interaction lists (“Old”). Precision values for “New” output, i.e. from the current pipeline, were chosen to match the Old precision values. CORUM version 2012 was used as a gold standard reference B. Run time for all modules on a non-performance PC using either the previously published version (“Old (2015)”, (Scott *et al.*, 2015)) or the current version (“New”).

### Predicting PPIs (Interactions.m)

Predicting protein-protein interactions (PPIs) is one of the primary functions of this pipeline. Figure 3 illustrates this process using a subset of D1 that contains ribosomal and proteasomal proteins. Each potential interaction, i.e. protein pair, is first identified as either a reference interaction (white), reference non-interaction, i.e. proteins in the reference that do not interact (black), or unknown (grey; Fig. 3A). To score each potential interaction, the similarity of each pair of co-fractionation profiles is then quantified using the five distance measures (Supp. Fig. 1; see Methods for definitions). Using these as input to the machine learning classifier, an interaction probability for each protein pair is then calculated, expressing how well each protein pair resembles the collection of reference PPIs (Fig. 3B).

**Figure 3:**
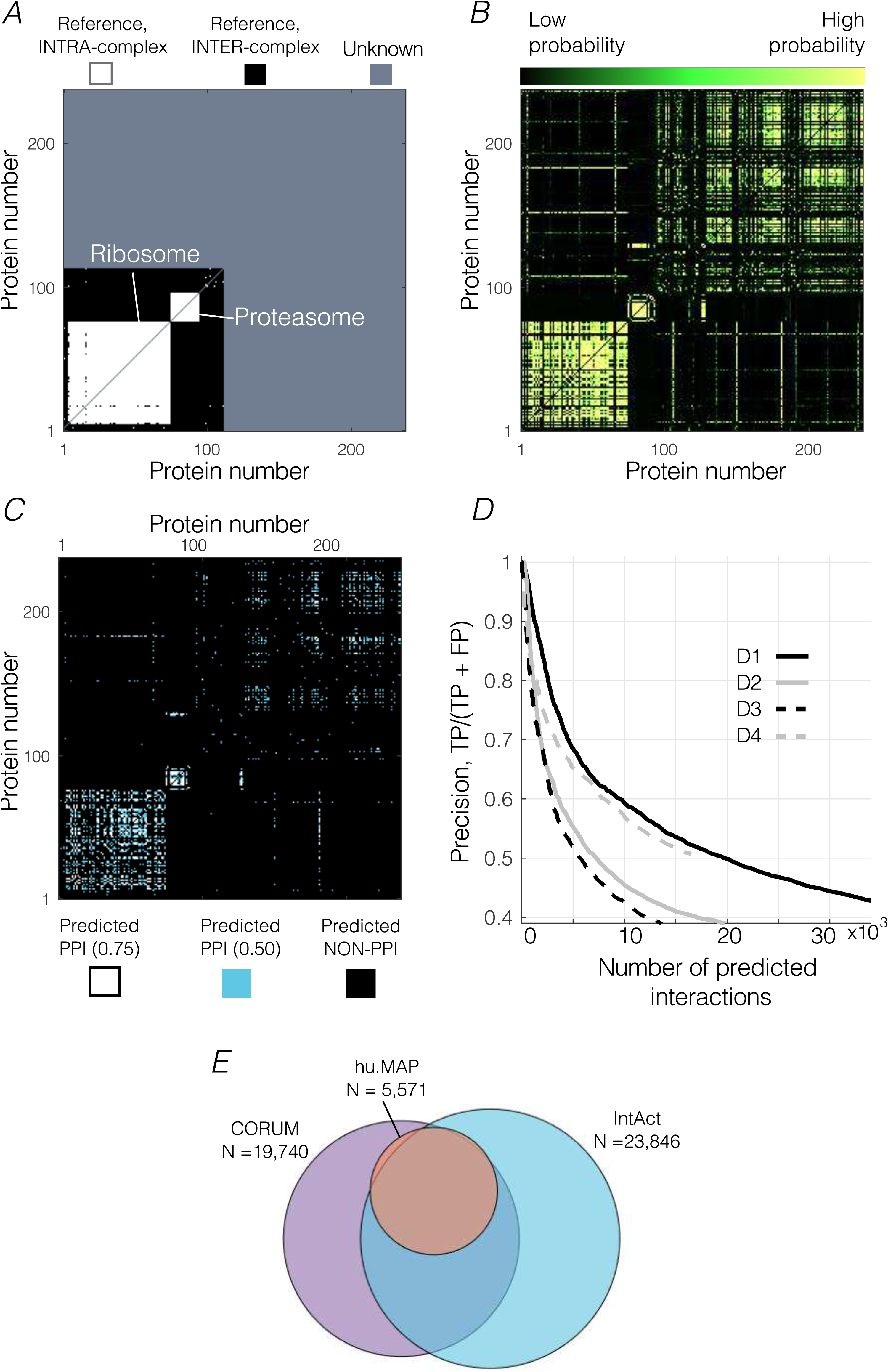
Predicting interactions (Interactions.m). A. Reference database. Subset of the CORUM reference database, including ribosomal and proteasomal proteins, expressed as a square pairwise matrix. Intra-complex interactions (white) are pairs of proteins from the same reference complex, inter-complex interactions (black) are pairs of proteins contained in the reference that are not co-complex members, and unknown/novel pairs (grey) have one or more protein not contained in the reference. Proteins are sorted according to their peak location. B. Interaction probability for each pair of proteins using the labels in A and distance measures. C. Square pairwise matrix of predicted interactions at two precision levels, 50% (0.50) and 75% (0.75). Interactions are predicted by applying a constant threshold to interaction score. D. Precision versus accumulated number of interactions. E. Predicted interactions using different gold standard references (CORUM, IntAct, and hu.MAP). 5,527 interactions were predicted from all three gold standards (overlap).

By applying a threshold to interaction probabilities outputted by the classifier, a final interaction list can be generated at a precision specified by the user. For example, a high confidence list containing an estimated 75% true positives (white), or a more inclusive, lower quality list with an estimated 50% true positives (cyan; Fig. 3C). In general, there is a tradeoff between quantity and quality when predicting PPIs, meaning that more PPIs can be predicted at the cost of lowering the quality, i.e. precision (Fig. 3D).

How does the number of quantified proteins affect the number of predicted interactions? To investigate, we analyzed random subsets of each dataset. Although there was considerable variability between datasets, in general there is an *N^2^* relationship between the number of proteins used as input to PrInCE and the number of interactions returned as output (Supp. Fig. 2). For all datasets, fewer than 500 quantified proteins resulted in less than 1,000 interaction at 50% precision. It is important to note that while PrInCE is designed to predict reference-like PPIs, it would be useless if it didn’t also predict *novel* interactions. That is, PrInCE must predict interactions that are not simply contained in the reference database. Indeed, for the subset of proteins shown in Figure 3 it can be seen that novel interactions are predicted (Fig. 3C, protein numbers 113 to 237). More broadly, all three datasets we used for benchmarking had thousands of novel PPIs predicted at 50% precision and hundreds to thousands of PPIs at 75% precision (Fig. 2A, Table 1). In particular, at 50% precision 16,019 interactions were predicted from D1 that are not contained in the reference.

Although PrInCE uses the CORUM database of protein complexes by default, in principal any suitable gold standard reference can be used. Since CORUM is potentially biased toward high stoichiometry complexes [24], we investigated the performance of PrInCE trained on two other gold standard protein complex databases: IntAct, a manually curated database of 1855 protein complexes [26], and hu.MAP, a database synthesized from three high throughput datasets totaling over 9,000 mass spectrometry experiments [27]. Using dataset D1 and predicting interactions at 50% precision, these databases produce 23,846 and 5,571 interactions, respectively (Fig. 3E). CORUM, which produces 19,740 interactions for D1 at the same precision level (Fig. 2A), is therefore comparable in performance to IntAct, both of which perform better than hu.MAP, possibly because CORUM and IntAct are hand curated. Therefore, the choice of gold standard reference is important, although all three gold standards lead to thousands of predicted interactions. Importantly, although the performance differs, a core set of interactions tends to be predicted regardless of the gold standard (Fig. 3E).

### Predicting protein complexes (Complexes.m)

Building on predicted PPIs, the second major output of PrInCE is protein complexes. Because buffer conditions in PCP-SILAC are relatively gentle on protein complexes, this module potentially identifies complexes that are unlikely to be identified by immunoprecipitation techniques. To do so, PPIs predicted by *Interactions.m* are weighted by their interaction score and input into the ClusterONE algorithm [16] to cluster individual PPIs into complexes.

Sorting co-fractionation profiles by their peak location (Fig. 4A) reveals the tendency for groups of proteins to co-elute (Fig. 4B). After analysis with PrInCE, some groups are predicted to be co-complex members. Figure 4C shows an example protein complex predicted by *Complexes.m*. The predicted complex (orange and purple) largely overlaps with the 20S proteasome contained in the CORUM reference database (black and purple). One novel member (P28065, orange) was predicted to be participating in the complex. Notably, while P28065 is not in the CORUM database, it is annotated as a proteasomal protein. Thus, using co-elution as the only source of evidence, PrInCE predicted one proteasomal protein as a novel co-complex member of the 20S proteasome.

**Figure 4:**
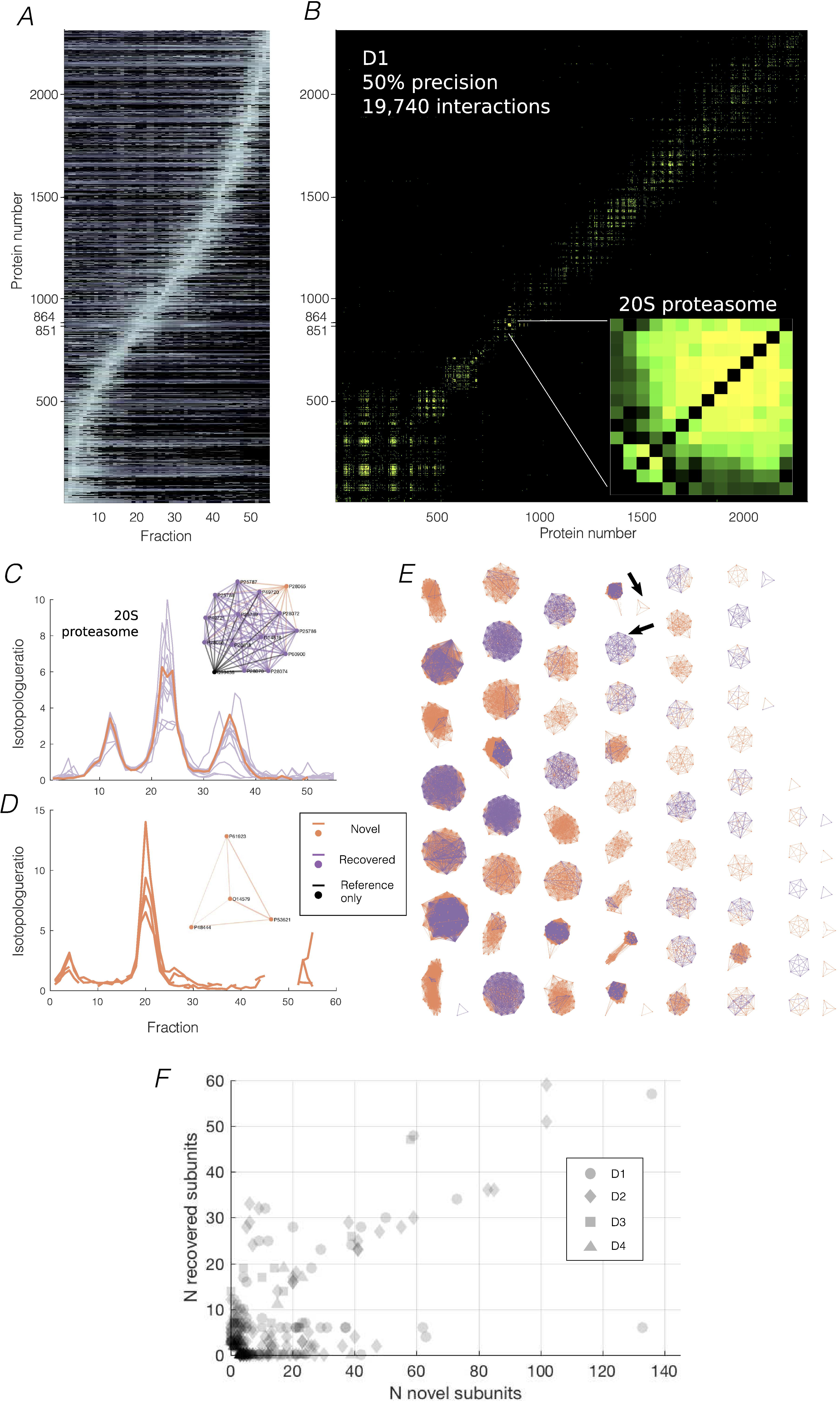
Predicting complexes (Complexes.m). A. 2,311 co-fractionation profiles from a single replicate of D1, sorted by peak location. Fourteen 20S proteasomal proteins group together (protein numbers 851-864). B. Square connection matrix for same proteins as A. Colour shows interaction score for all 19,740 interactions with score greater than 0.50. Inset: Close up of the 14x14 connection matrix for 20S proteasomal members plus other proteins (protein numbers 851-865). C. Co-fractionation profiles for the 14 proteins from B inset, which also correspond to a predicted complex. Profiles of complex members (left) all have a similar shape. When compared to its closest match in CORUM, the 20S proteasome, this predicted complex had 13 overlapping proteins (purple), as well as one protein in the predicted complex that was not in the 20S proteasome (orange). Additionally, there was a single protein from the 20S proteasome that was not in the predicted complex (black). D. Example predicted complex with no match in the CORUM database. E. Force diagrams for all 71 predicted complexes from 19,740 interactions in D1. Same colouring scheme as D and E. Proteins in known complexes that were not predicted (i.e. Reference-only, black) are omitted for clarity. F. Predicted complexes are composed of known (“recovered”) subunits and novel subunits. Data is from all four datasets. The size of each predicted complex is the sum of novel and recovered members.

PrInCE is also capable of predicting entirely novel protein complexes. For example, a four member complex was predicted in dataset D1, of which no proteins were in CORUM (Fig. 4D). Reassuringly, these four proteins (P61923, P53621, P48444, O14579) are all subunits of the coatomer protein complex, a known complex that, while not present in the CORUM database, has substantial low throughput [28–30] and high throughput evidence [6, 8, 15] supporting its existence. For all complexes predicted by the pipeline (e.g. Fig. 4E; D1, 71 complexes, median size 14), each complex predicted by ClusterONE is matched to a reference complex when possible. Of the 71 protein complexes predicted for D1, 20 were entirely novel, i.e. had no matching reference complex. In general, PrInCE predicts both entirely novel protein complexes and those that recover existing complexes while predicting novel members. The four datasets analyzed in this study produced a total of 291 protein complexes, of which 169 were at least partially matched to a CORUM complex. On average, 31% of complex subunits were recovered from known complexes while the remaining were novel subunits (Fig. 4F).

### Validation of predicted interactions and complexes

No method for determining protein interactions is perfect, and higher-throughput methods tend to recover noise along with biologically meaningful signal. We estimate how much noise is in the final interaction list by comparing it to a reference of known interactions, e.g. CORUM, and quantifying the signal to noise ratio in terms of precision, i.e. *TP/*(*TP* + *FP*). In order to validate that we are separating signal from noise in a biologically meaningful way, we sought to establish the biological significance of interaction lists generated by PRInCE using independent evidence. First, we wanted to confirm that the measure we use to rank the quality of predicted interactions, interaction score, is a useful way to identify which interactions are more likely to be true positives. To do so, we tested whether proteins in high score PPIs are more likely to share annotation terms than low score interactions. Indeed, for every GO-slim annotation category, as interaction score increased, so did the proportion of interactions sharing at least one annotation term (Fig. 5A, Supp. Table 1). Similarly, interacting protein pairs were more likely to be significantly coexpressed across human tissues (Pearson correlation coefficient ≥ 0.75) (Fig. 5B), share at least one subcellular localization term (Supp. Fig. 3A), and have a structurally resolved domain-domain interaction (Supp. Fig. 3B). Therefore, the ranking system used by this pipeline is biologically significant, as demonstrated by independent sources of evidence.

**Figure 5:**
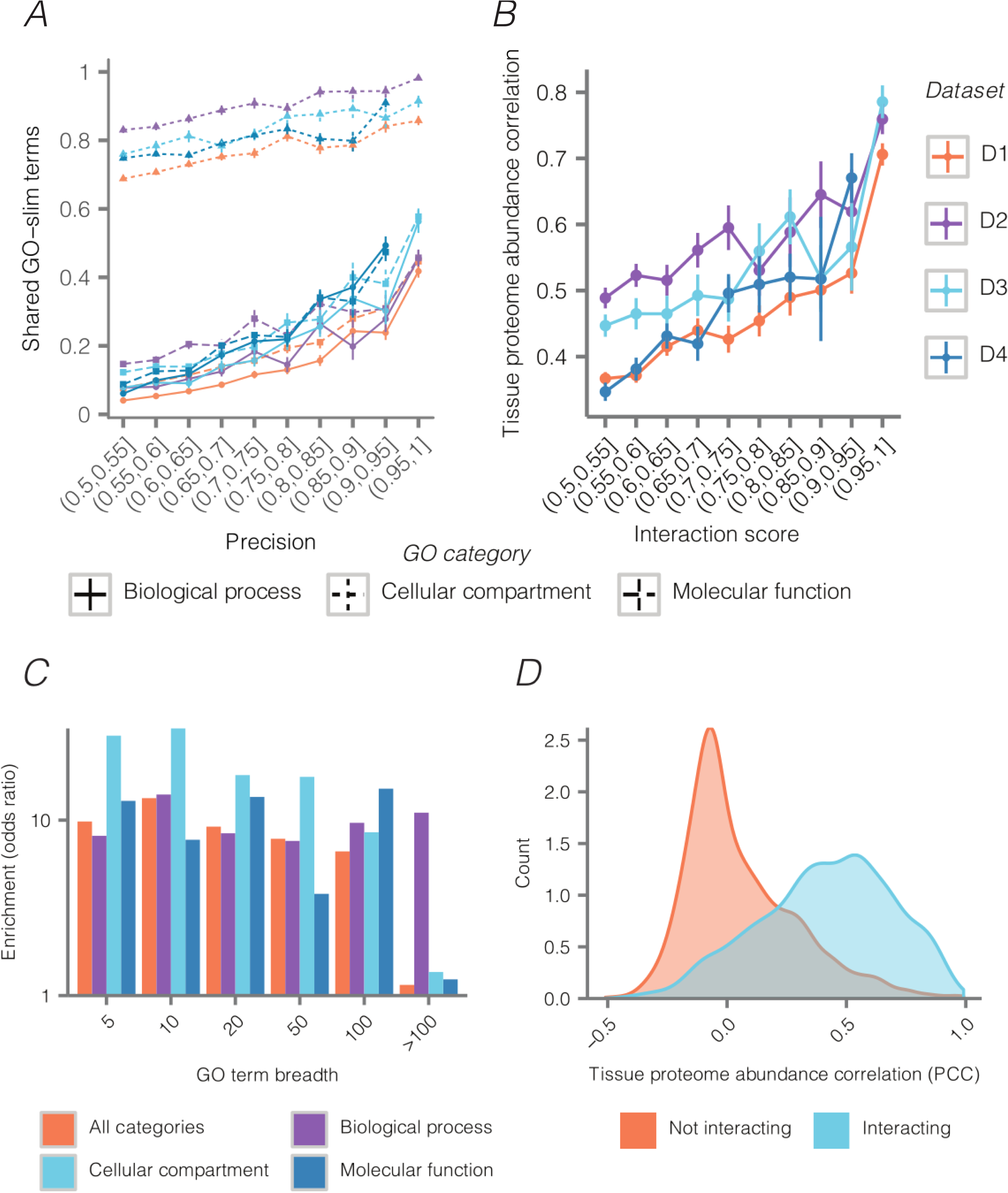
Predicted interactions are enriched for biologically significant attributes, and the degree of enrichment reflects interaction score. A. Fraction of interacting proteins with at least one shared GO-slim term as a function of interaction score and ontological domain. Triangle: biological process. Square: cellular component. Circle: molecular function. B. Tissue proteome abundance [21] correlation (Pearson correlation coefficient) as a function of interaction score. C. Interacting proteins in the apoptosis dataset are enriched for shared GO-slim terms relative to non-interacting protein pairs at diverse GO term breadths. D. Distribution of tissue proteome abundance correlations (Pearson correlation coefficients) for interacting and non-interacting protein pairs in D1.

How do predicted interactions differ from predicted non-interactions? A well-performing pipeline returns predicted classes that are, at least by some measures, cleanly separated. To assess this, we first compared Jaccard indices [24], which measure the degree to which protein pairs share annotation terms, between non-interacting protein pairs (cyan), medium-confidence predictions (orange), and high-confidence (purple; Supp. Fig. 3C, Supp. Fig. 4A-C). Compared to non-interacting proteins, high-confidence interactions show a bias towards larger Jaccard indices, as do medium-confidence interactions, although to a lesser degree.

We next used enrichment values to quantify the tendency for predicted interacting proteins to share annotation terms. In general, interacting proteins were about 10x more likely to share GO annotation terms than non-interacting proteins (Fig. 5C, Supp. Fig. 4D-F). Moreover, enrichment was relatively independent of the breadth of the annotation terms, where breadth describes the number of annotated proteins per annotation term [31]. We found that interacting proteins were significantly enriched for nearly all validation measures used here (Table 2). Finally, comparing how well tissue-dependent protein abundance correlates between protein pairs [21] shows that protein abundance correlates significantly better between predicted interacting protein pairs versus predicted non-interactions (Fig. 5D, Supp. Fig. 4G-J). Therefore, predicted interactions returned by PrInCE are significantly higher quality than predicted non-interactions, as supported by measures which are independent of the evidence used within the pipeline. The same analysis was repeated to compare interactions predicted by PrInCE to previously published interaction lists [8, 11]. To do so, we matched the number of interactions in the published lists by taking that number of top-ranked interactions predicted by PrInCE. In 15 out 18 comparisons of enrichment values, interactions predicted by PrInCE were measured to be higher quality than previously published lists (Supp. Table 2).

**Table 2.**
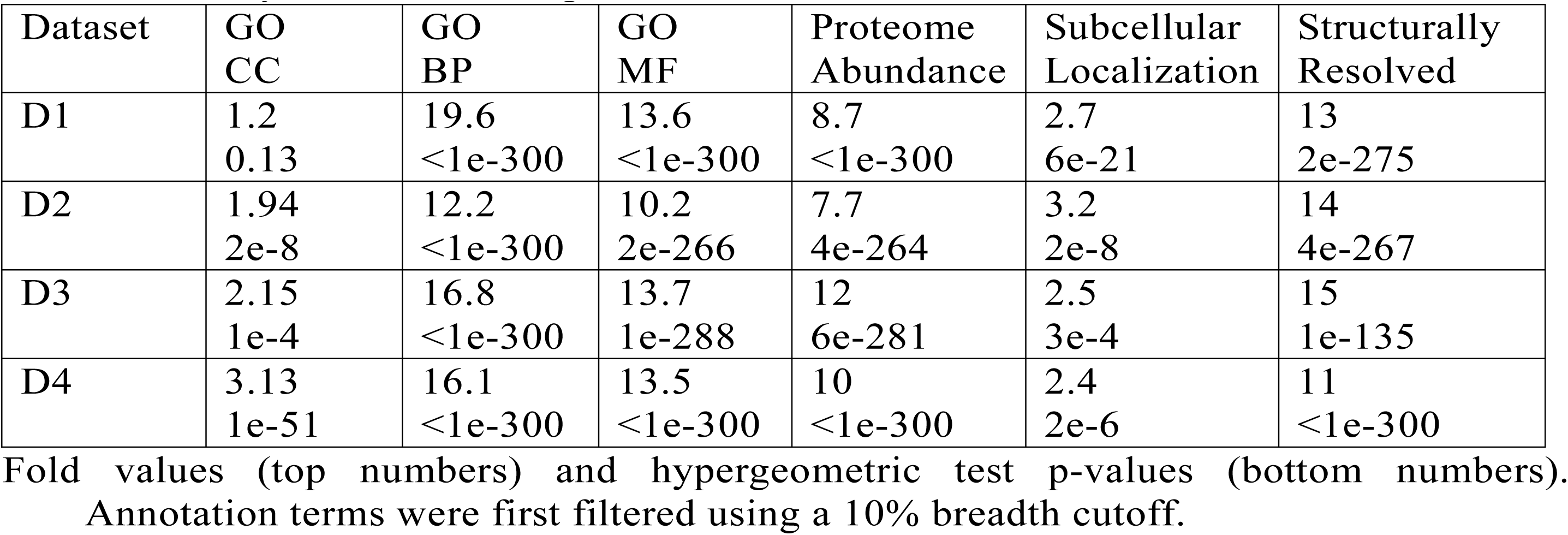
Interacting versus non-interacting terms for shared annotation terms (GO, Subcellular Localization), tissue-dependent proteome abundance, and shared structurally resolved binding domains.

| Dataset | GO<br>CC | GO<br>BP | GO<br>MF | Proteome<br>Abundance | Subcellular<br>Localization | Structurally<br>Resolved |
| --- | --- | --- | --- | --- | --- | --- |
| D1 | 1.2<br>0.13 | 19.6<br><1e-300 | 13.6<br><1e-300 | 8.7<br><1e-300 | 2.7<br>6e-21 | 13<br>2e-275 |
| D2 | 1.94<br>2e-8 | 12.2<br><1e-300 | 10.2<br>2e-266 | 7.7<br>4e-264 | 3.2<br>2e-8 | 14<br>4e-267 |
| D3 | 2.15<br>1e-4 | 16.8<br><1e-300 | 13.7<br>1e-288 | 12<br>6e-281 | 2.5<br>3e-4 | 15<br>1e-135 |
| D4 | 3.13<br>1e-51 | 16.1<br><1e-300 | 13.5<br><1e-300 | 10<br><1e-300 | 2.4<br>2e-6 | 11<br><1e-300 |

Fold values (top numbers) and hypergeometric test p-values (bottom numbers). Annotation terms were first filtered using a 10% breadth cutoff.

Calculating the precision of the interactions predicted by PrInCE is crucial for minimizing the number of false positives. To estimate precision, both the numbers of true and false positives must be calculated. The reference database provides a list of true positive interactions (intra-complex). However, since no comparable database of false positive interactions exists, we make the assumption that pairs of interacting proteins which are both present in the reference, but not reported by the reference to interact, are false positives (inter-complex). Several of these false positives are likely to be true interactions that simply have not been previously discovered and thus not included in the reference, meaning that PrInCE likely underestimates the true precision of the interactions. Using the method outlined in [25] to re-estimate precision, we found that, indeed, the stated precision is a conservative estimate of the quality of the predicted interaction list (Fig. 6).

**Figure 6:**
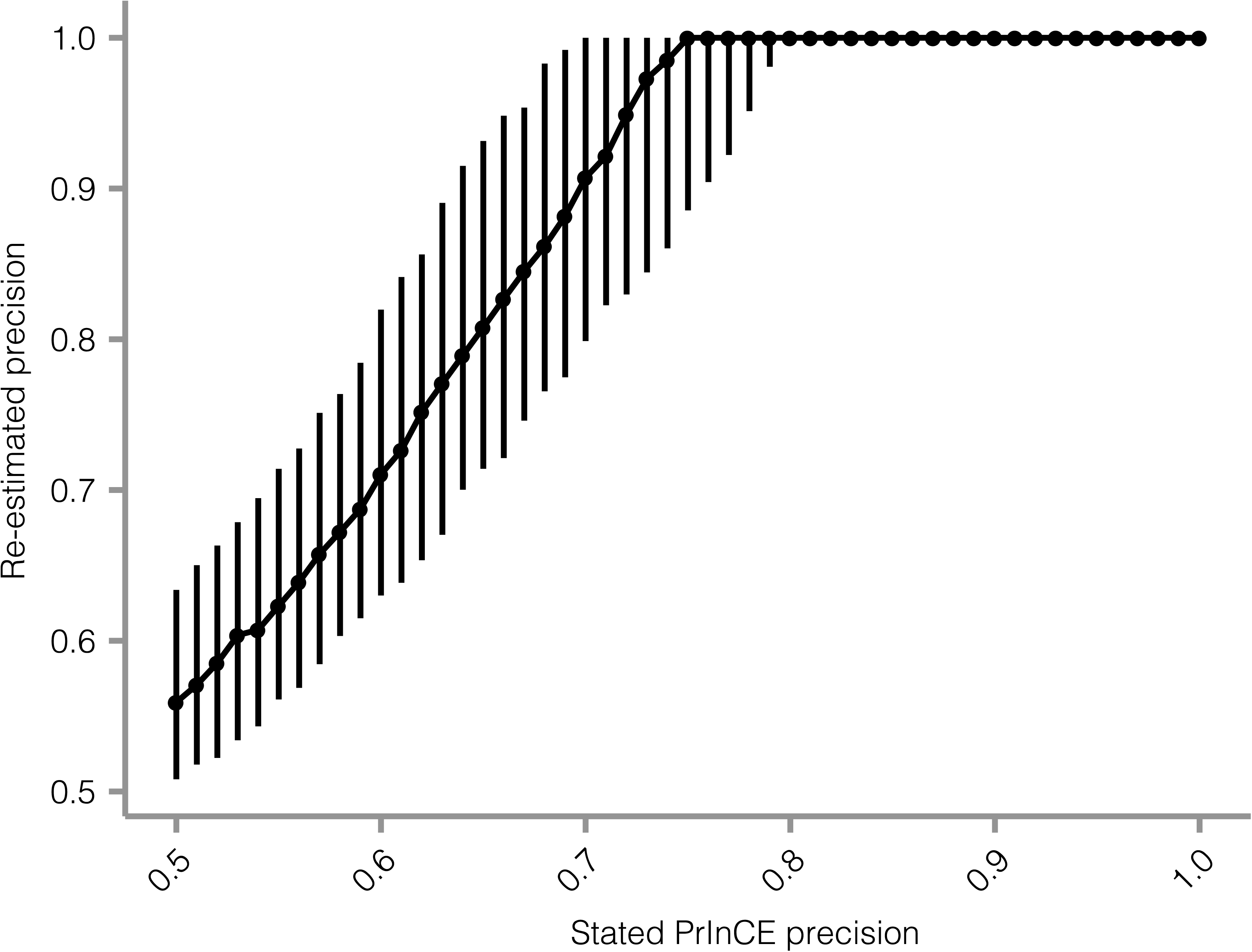
PrInCE precision of the predicted interaction list is a conservative estimate of the number of false positives. Predicted interaction lists were generated for dataset D1 at multiple user-defined precision levels (PrInCE precision), and their precision was re-estimated (Mrowka precision [25]). PrInCE lists were generated using a random 2/3 subset of the CORUM reference and precision was re-estimated using the remaining 1/3. Median values from 100 iterations are shown, and bars show the interquartile range.

Finally, we explored the topological properties of the predicted network, i.e. how the network is connected. Specifically, as is postulated for other PPI networks returned by high-throughput techniques [24], we validated the hypothesis that predicted networks should consist of small subsets of highly connected proteins, which are more loosely linked to each other by relatively weak connections. This connectivity structure denotes well-defined subgraphs connected by weaker signaling and/or spurious false positive interactions. To analyze the topology, we used an approach described by [24], wherein interactions are removed sequentially from the network: removing the lowest confidence interactions first should *fragment* the network by revealing islands of isolated subgraphs; removing the highest confidence interactions should lead to no fragmentation. Indeed, removing low confidence interactions first produced a network with a greater number (Supp. Fig. 5A, purple) of relatively smaller subgraphs (Supp. Fig. 5B), i.e. fragmentation. Removing interactions in this order rapidly fragmented the largest subgraph (Supp. Fig. 5C). Removing high-confidence interactions first did not have this effect (Supp. Fig. 5, orange). Similar results were obtained for other datasets (Supp. Fig. 5E-P).

## 4 Discussion

A machine learning classifier provides improvements over simply sorting protein-protein pairs by how similarly they co-elute, as it provides an automated method for combining multiple measures of co-elution. We chose the Naive Bayes classifier because it is computationally inexpensive and surprisingly powerful given its relative simplicity. Indeed, when comparing the Naive Bayes (“fitcnb”, Matlab) to a Support Vector Machine classifier (“fitcsvm”, Matlab) we found the Naive Bayes performed on par or better with respect to the number of predicted interactions at a given precision level (data not shown).

One limitation of our technique is that it requires a suitable gold standard reference of known protein complexes. Since protein complexes are variable, not all known interactions will occur at any one time or under one set of biological conditions. Therefore, the suitability of a reference database, determined by the fraction of gold standard interactions that were indeed physically interacting in the sample, is crucial. Failure of the data to adhere to the reference will result in poor classification performance and, ultimately, a short or empty list of predicted interactions. For mammalian experiments, we preferentially use the CORUM database as our reference, although thousands of interactions were predicted with both the IntAct [26] and hu.MAP [27] gold standards (Fig. 3E). It is also possible to make a custom reference list of complexes.

Early versions of this pipeline were designed for the analysis of (PCP-) SILAC datasets. A major strength of SILAC experiments is that they allow conditional experiments to be performed simultaneously, minimizing experimental variability between conditions. However, the analysis here of dataset D3, a surrogate for a non-SILAC labelled dataset, demonstrates that PrInCE is not limited to analyzing SILAC data. In fact, PrInCE can analyze any dataset with co-fractionation profiles for single proteins where co-fractionation is meaningful evidence of co-complex membership, and for which there exists a suitable reference.

## 5 Conclusions

PrInCE provides a powerful, easy to use tool for predicting interactomes from co-fractionation experiments. Building on preliminary versions of a bioinformatics treatment [8, 11], PrInCE predicts nearly twice as many protein interactions at the same stringency with a 97% decrease in run time (Fig. 2). PrInCE also offers increased functionality over previous versions, providing a module for automated, optimized prediction of protein complexes using the ClusterONE algorithm [16]. Importantly, PrInCE is available as a standalone executable program, meaning access to Matlab is not required. Finally, at the same number of interactions, interactions predicted by PrInCE are higher quality than previous versions, as quantified by a greater enrichment of shared annotation terms (Supp. Table 2).

## 5 Declarations

### Ethics approval and consent to participate

Not applicable.

### Consent for publication

Not applicable.

### Availability of data and material

The three published co-elution datasets analyzed during the current study are available from doi: 10.1038/nmeth.2131, doi: 10.1016/j.jprot.2014.10.024, and doi: 10.15252/msb.20167067.

## Authors’ contribution

RGS analyzed and interpreted the data, wrote the PrInCE software, and drafted the manuscript and revised it. MS performed the validation analysis and critically revised the manuscript. NS made significant contributions to conception and design of the study and acquired two of the datasets analyzed here. LF made substantial contributions to conception and design of the study and critically revised the manuscript. All authors read and approved the final manuscript.

## Acknowledgements

We thank A. Prudova and A. McAfee for critical suggestions. M.A.S. is supported by a CIHR Frederick Banting and Charles Best Canada Graduate Scholarship, a UBC Four Year Fellowship, and a Vancouver Coastal Health-CIHR-UBC MD/PhD Studentship Award.

